# Polyfunctional IL-17A+ MAIT cells are expanded in the peripheral blood of patients with HLA-B27+ axial spondyloarthritis

**DOI:** 10.1101/2022.01.17.475829

**Authors:** Micah Lefton, Nihaarika Sharma, Akash R. Patel, Jeffrey A. Sparks, Joerg Ermann

## Abstract

**Objectives:** Studies in axial spondyloarthritis (axSpA) have yielded conflicting results regarding the identity of the major IL-17A-producing lymphocyte populations. The goal of this study was to comprehensively assess the production of IL-17A and related cytokines by peripheral blood lymphocytes in axSpA.

**Methods:** Peripheral blood mononuclear cells were isolated from patients with axSpA and healthy controls matched for age, sex and HLA-B27 status. Unstimulated cells and cells activated with PMA/Ionomycin were analyzed by 25-parameter fluorescent flow cytometry. Data were analyzed by hierarchical gating, UMAP and SPICE.

**Results:** Except for a reduced frequency of mucosal-associated invariant T (MAIT) cells and natural killer (NK) cells, there were no other significant differences in abundance of major lymphocyte populations in axSpA patients compared with controls. Increased IL-17A production in axSpA was observed in total non-B lymphocytes and in MAIT cells. The fraction of MAIT cells expressing the tissue residency markers CD69 and CD103 was increased in axSpA. CD103 positive MAIT cells were enriched for IL-17A producers. axSpA patients demonstrated an expansion of MAIT cell subsets producing IL-17A, IL-17F, GM-CSF and TNF. This expansion was only observed in HLA-B27+ patients.

**Conclusions:** We document an expansion of polyfunctional IL-17A+ MAIT cells in the peripheral blood of HLA-B27+ patients with axSpA. These results are consistent with the implied role of intestinal dysbiosis or inflammation in axSpA pathogenesis.

**Key messages:** What is already known about this subject?

- Various IL-17A-producing lymphocyte populations have been implicated in the pathogenesis of axSpA.

What does this study add?

- Polyfunctional MAIT cells capable of producing IL-17A, IL-17F, GM-CSF and TNF are expanded in the peripheral blood of HLA-B27+ patients with axSpA.

How might this impact on clinical practice or future developments?

- Overproduction of IL-17A by MAIT cells is the most consistent finding of peripheral blood lymphocyte studies in axSpA.
- Our data support the pathogenetic link between intestinal and axial inflammation in axSpA.

## Introduction

Axial spondyloarthritis (axSpA) including its more severe variant ankylosing spondylitis (AS) is an immune-mediated disease predominantly of the axial skeleton.[1] Multiple lines of evidence point toward a central role for lymphocytes in the pathogenesis of the disease. This includes the strong association with the MHC class I allele HLA-B27 as well as weaker associations, mostly identified in genome-wide association studies, with genes that encode transcription factors, cytokines and signaling modules involved in lymphocyte development, differentiation and function.[2] Antibody-mediated inhibition of the lymphocyte-derived cytokine IL-17A is now an established therapy for axSpA.[3, 4]

Studies in AS and axSpA have yielded conflicting results regarding the identity of the IL-17A-producing cells.[5, 6] Given the difficulties in obtaining tissue for analysis in axSpA, these studies have been almost exclusively performed in peripheral blood. Early studies focused on CD4+ T cells and described an expansion of IL-17A-producing Th17 cells.[7, 8]. More recently, increased IL-17A production in axSpA was reported for a variety of innate-like lymphocytes including γδ T cells [9], invariant natural killer T (iNKT) cells [10] and mucosal-associated invariant T (MAIT) cells.[11–13] A comprehensive analysis of these cell types in the same study has not been performed. It is therefore not clear whether there is a general dysregulation of IL-17A production in axSpA or whether subsets of patients can be distinguished with selective expansion of distinct IL-17A+ lymphocyte populations.

To address this question, we developed a high-dimensional flow cytometry panel for the comprehensive analysis of cell surface markers and intracellular cytokines. A cohort of patients with axSpA was compared to healthy controls matched for age, sex and HLA-B27 status. Consistent with previous reports, we found a reduction of total MAIT cells in axSpA peripheral blood associated with a higher frequency of IL-17A positive cells, many of which produced additional cytokines such IL-17F, GM-CSF and TNF. The expansion of these polyfunctional MAIT cells was only seen in HLA-B27+ patients.

## Methods

### Subject Recruitment

Patients with axial spondyloarthritis (axSpA) were recruited in the Arthritis Center at Brigham and Women’s Hospital (BWH), Boston, MA. The HLA-B27 status of the axSpA patients was retrieved from medical records. Healthy controls were recruited from the Mass General Brigham (MGB) Biobank, a research repository at BWH, Massachusetts General Hospital and affiliated sites in the Greater Boston, MA, area.[14, 15] Participants in MGB Biobank provide blood to be stored for research purposes, allow for access to their electronic health record and agree to be re-contacted for other research studies. About one-third of participants were randomly selected to be genotyped using ILUMINA arrays. We identified HLA-B27+/− individuals using SNP2HLA [16] and applied a filter to screen out subjects with SpA-related diseases and other inflammatory conditions or chronic conditions that might affect immune status. We then invited a random sample of 500 HLA-B27+/− healthy controls less than 55 years of age to provide a peripheral blood sample specifically for this study enrolling a total of 32 subjects that were verified to be healthy on medical record review. Prior to the flow cytometry experiment, we matched available axSpA and healthy control specimens for age, sex and HLA-B27 status selecting n=28 healthy controls and n=28 axSpA samples for inclusion in the study (Suppl. Figure S1A). All subjects gave informed consent. The study was approved by the MGB IRB. Patients and the public were not involved in the design, or conduct, or reporting, or dissemination plans of this research.

### PBMC Isolation and Storage

Peripheral blood was obtained by venous blood draw. Peripheral blood mononuclear cells (PBMC) were isolated using Ficoll-Paque PLUS (GE Healthcare) and cryopreserved in freezing medium containing fetal bovine serum (FBS) (Gemini) and 10% dimethyl sulfoxide (DMSO) (Sigma Aldrich).

### *In Vitro* Stimulation

Thawed PBMCs were seeded in 96-well round bottom plates at 2 x 10^6^ cells/well in Iscove’s Modified Dulbecco’s Medium (Gibco) supplemented with 10% fetal bovine serum (Sigma Aldrich), sodium pyruvate (Cellgro), penicillin and streptomycin (Sigma Aldrich), nonessential amino acids (Cellgro) and 2-mercaptoethanol (Gibco). Cells were stimulated with 50 ng/mL PMA plus 100 ng/mL Ionomycin (Cell Stimulation Cocktail, Thermo Fisher Scientific), in the presence of monensin (2 ug/ml, GolgiStop, BD Bioscience) and brefeldin A (1 ug/ml, GolgiPlug, BD Bioscience) for 6 hours.

### Flow Cytometry

Unstimulated PBMCs were stained using the antibodies in Suppl. Table 1. Stimulated cells were first stained for surface markers. After fixation in IC Fixation Buffer (Thermo Fisher Scientific), intracellular staining for cytokines was performed in Permeabilization Buffer (Thermo Fisher Scientific). Suppl. Table 2 lists the antibodies used for staining stimulated PBMCs. Cells were analyzed on a FACSymphony flow cytometer (BD Bioscience) and data were analyzed using FlowJo v10.8.0 (BD Bioscience). Multi-cytokine analysis was performed using SPICE v6.[17] See Suppl. Figure S1B for workflow and Suppl. methods for additional detail.

### Statistical Analysis

Statistical analysis and graphing were done using Prism v9.0.2 (Graphpad Software) and R v4.1.0.[18] Student t test or ANOVA were used for the analysis of parametric data. Mann Whitney U test or Kruskal-Wallis test were used for non-parametric data. A p-value <0.05 was considered statistically significant.

## Results

### Reduced frequency of MAIT and NK cells in axSpA

To characterize lymphocyte populations in axSpA, we developed two 25-parameter fluorescent flow cytometry panels (Suppl. Table 1 and 2). We applied these panels to a collection of PBMC specimens from patients with axSpA and healthy controls matched for age, sex and HLA-B27 status (Table 1, Suppl. Figure 1A).

**Table 1.** Study subjects.

|  | Axial Spondyloarthritis (axSpA)<br>n = 28 | Healthy Controls<br>n = 28 |
| --- | --- | --- |
| Age (years), median (range) | 42 (26 – 57) | 42.5 (24 – 55) |
| Males, n (%) | 17 (61%) | 15 (54%) |
| HLA-B27+, n (%) | 19 (68%) | 15 (54%) |
| AS, n (%) | 16 (57%) | N/A |
| CRP (mg/L), median (range) | 2.9 (0.4 – 31.5) | N/A |
| BASDAI, median (range) | 2.9 (0.4 – 10) | N/A |
| NSAID, n (%) | 17 (60%) | 5 (18%) |
| Anti-TNF, n (%) | 17 (60%) | 0 (0%) |
| Anti-IL-17A, n (%) | 2 (7%) | 0 (0%) |
| cDMARD, n (%) | 0 (0%) | 0 (0%) |

We used hierarchical gating (Suppl. Figure 2) to identify CD4+ T cells, CD8+ T cells, DN αβ T cells, MAIT cells, γδ T cells distinguished as Vδ1, Vδ2 and other γδ T cells, NK cells and innate lymphoid cells (ILCs) in unstimulated PBMCs. The frequencies of these populations were similar in healthy controls and axSpA patients with two exceptions (Figure 1). Consistent with reports by others, we found that the frequency of MAIT cells (identified here as CD3+αβTCR+MR-1 5-OP-RU tetramer+ cells) was significantly reduced in axSpA patients.[11–13] The frequency of NK cells (CD3-CD56+) was also lower in axSpA than in healthy controls. There were no differences in the frequency of γδ T cells or any of the other major lymphocyte populations. We also did not find differences between axSpA patients and healthy controls (Suppl. Figure 3) in CD4+ or CD8+ T cell subsets distinguished by CD45RA and CCR7 expression, Treg cells (CD45RA-CD127-CD25+) or CD8+ InEx cells (CD45RA-CD49a+CD103+).[19]

**Figure 1.**
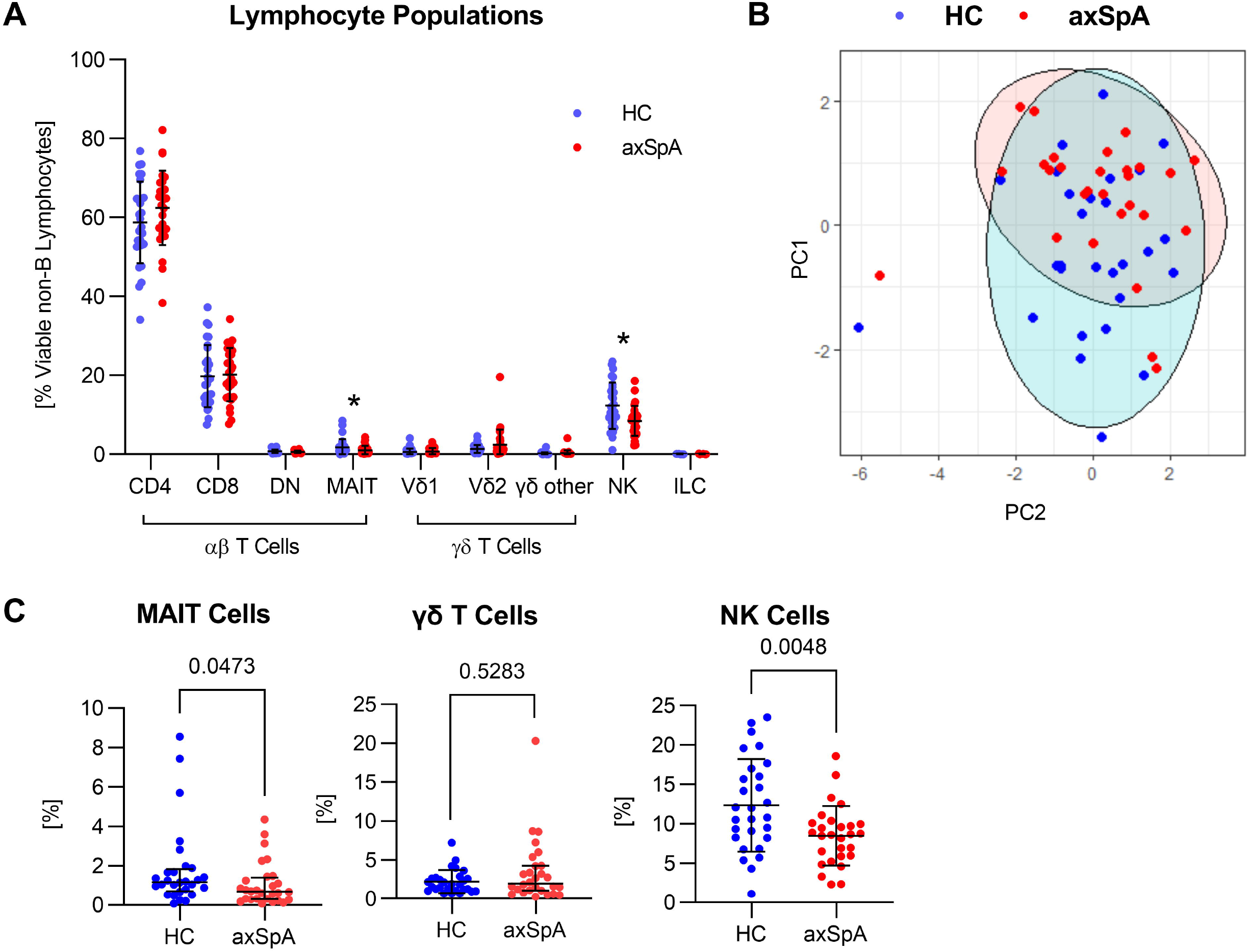
Decreased frequency of MAIT and NK cells in axSpA patients. **A.** Frequencies of major lymphocyte populations in patients with axSpA (n=28) and healthy controls (n=28) determined by hierarchical gating. Numbers are the percentage of viable non-B lymphocytes. Bars represent means and standard deviation. Asterisks indicate a p value < 0.05. **B.** PCA analysis of lymphocyte population frequencies: axSpA and healthy control clusters are largely overlapping. **C**. “Magnification” of data from panel A for MAIT cells, total γδ T cells (sum of Vδ1, Vδ2 and other γδ T cells) and NK cells. The frequency of MAIT cells (p=0.0473) and NK cells (p=0.0048) was reduced in axSpA subjects. No difference in the frequency of total γδ T cells (p=0.5283).

### Increased frequency of IL-17A+ and IL-17F+ MAIT cells in axSpA

A second aliquot of cells was stimulated with PMA/Ionomycin and analyzed for production of IL-17A, IL-17F, IL-22, GM-CSF, IFNγ and TNF by intracellular staining (Figure 2A). The frequency of IL-17A+ total non-B lymphocytes was significantly higher in axSpA patients than in healthy controls (Figure 2B). As a first pass analysis, the expression of individual cytokines was examined in the major lymphocyte populations identified by hierarchical gating. Summary data in Suppl. Figure 4 show that the pattern of cytokine expression differed substantially between cell types. Differences between axSpA patients and controls were minor and limited to expression of IL-17A and IL-17F. MAIT cells from axSpA patients had a higher fraction of IL-17A+ cells than healthy controls. CD4+ and CD8+ T cells also showed a trend for increased IL-17A production in axSpA, but this was not statistically significant. There was no difference in IL-17A expression by total γδ T cells or γδ T cell subsets from axSpA patients and controls (Figure 2C).[9] IL-17F, which is structurally and functionally similar to IL-17A, could only be detected in CD4+ T cells and MAIT cells. AxSpA MAIT cells had a higher frequency of IL-17F positive cells than healthy controls, while there was no difference in CD4+ T cells (Figure 2D). IL-17F expression was largely limited to cells that expressed IL-17A. The frequency of IL-17A/IL-17F double positive cells was higher in MAIT cells from axSpA patient compared to controls, there was a non-significant trend in CD4+ T cells from axSpA patients (Figure 2E). We did not find differences in expression of IL-22, GM-CSF, IFNγ or TNF when analyzed individually in any of the nine major lymphocyte populations (Suppl. Figure 4).

**Figure 2.**
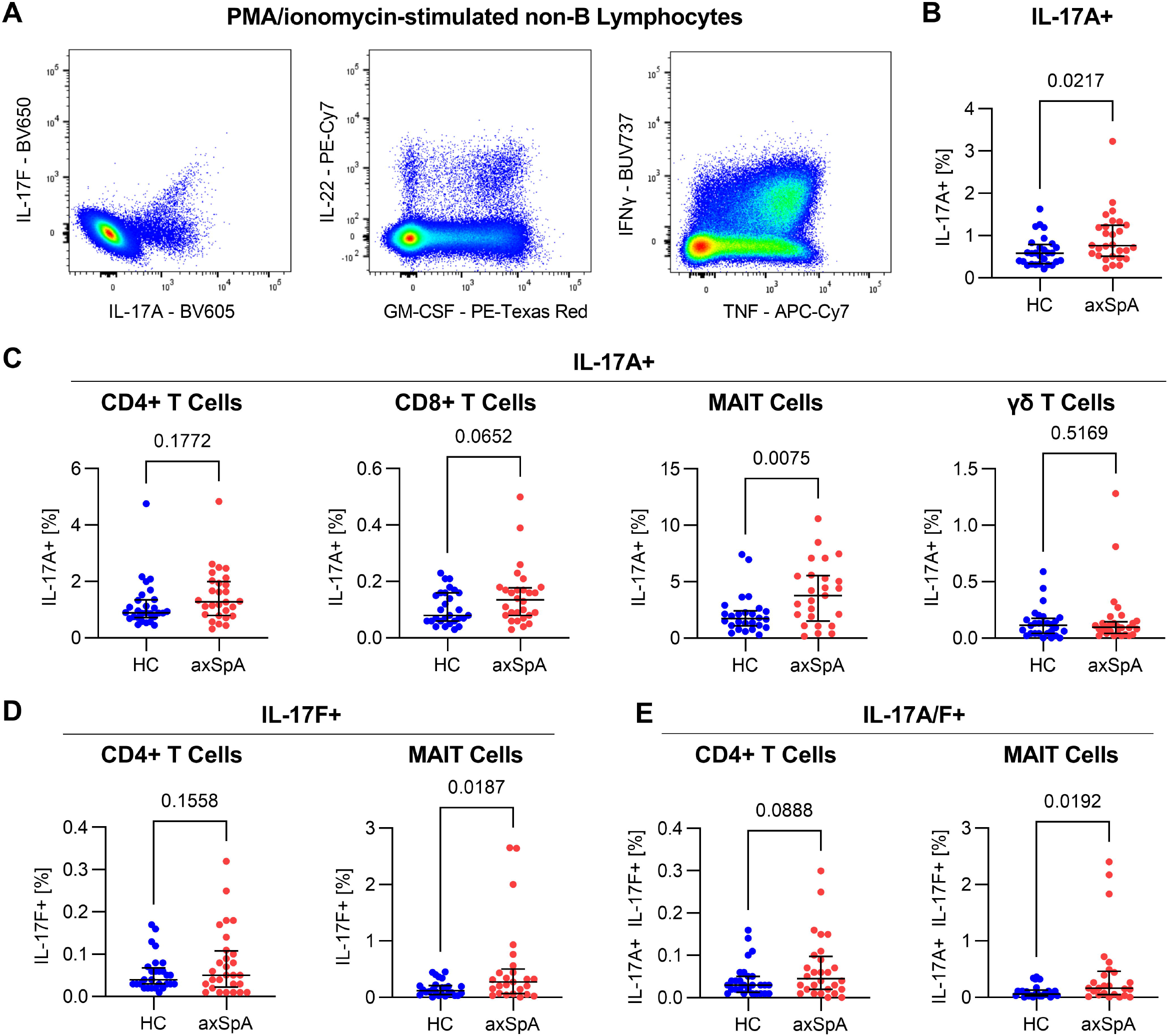
Expansion of IL-17A+ and IL-17F+ MAIT cells in axSpA patients. PBMCs were stimulated with PMA/ionomycin for 6 hours. **A**. Example plots for intracellular staining of IL-17A, IL-17F, IL-22, GM-CSF, IFNγ and TNF in total non-B lymphocytes. **B**. The IL-17A+ fraction of total non-B lymphocytes was higher in axSpA patients than in healthy controls (p=0.0217). **C**. There was a trend for increased fractions of IL-17A+ cells in CD4+ and CD8+ T cells in axSpA while axSpA patients had significantly more IL-17A+ MAIT cells (p=0.0075). No difference in the frequency of IL-17A+ γδ T cells. **D**. IL-17F positivity trended higher in CD4+ T cells from axSpA patients and was significantly increased in MAIT cells (p=0.0187). **E**. Similarly, IL-17A/IL-17F double positive cells trended higher CD4+ T cells and were significantly increased in MAIT cells from axSpA patients.

### Polyfunctional MAIT cells are expanded in axSpA

Considering the significant differences in expression of IL-17A and IL-17F in MAIT cells (but not other cell types) from axSpA patients and healthy controls, we focused subsequent analyses on this cell population. As expected for peripheral blood MAIT cells in adults, the MAIT cells in axSpA patients and controls were CD161+, CD127+ (IL-7R+) and largely CD8+, CD45RA- and CCR7-(Suppl. Figure 5).[20, 21] MAIT cells at mucosal sites strongly express the tissue residency markers CD69 and CD103.[22, 23] These markers are also expressed on some circulating MAIT cells.[24] Comparing MAIT cells from axSpA patients and healthy controls, we found an increased fraction of CD69-CD103+ cells in the patients, CD69+CD103- and CD69+CD103+ fractions also trended higher in this group (Figure 3A). IL-17A production upon PMA/Ionomycin stimulation was enriched in CD69-CD103+ cells, both in axSpA patients and healthy controls (Figure 3B).

**Figure 3.**
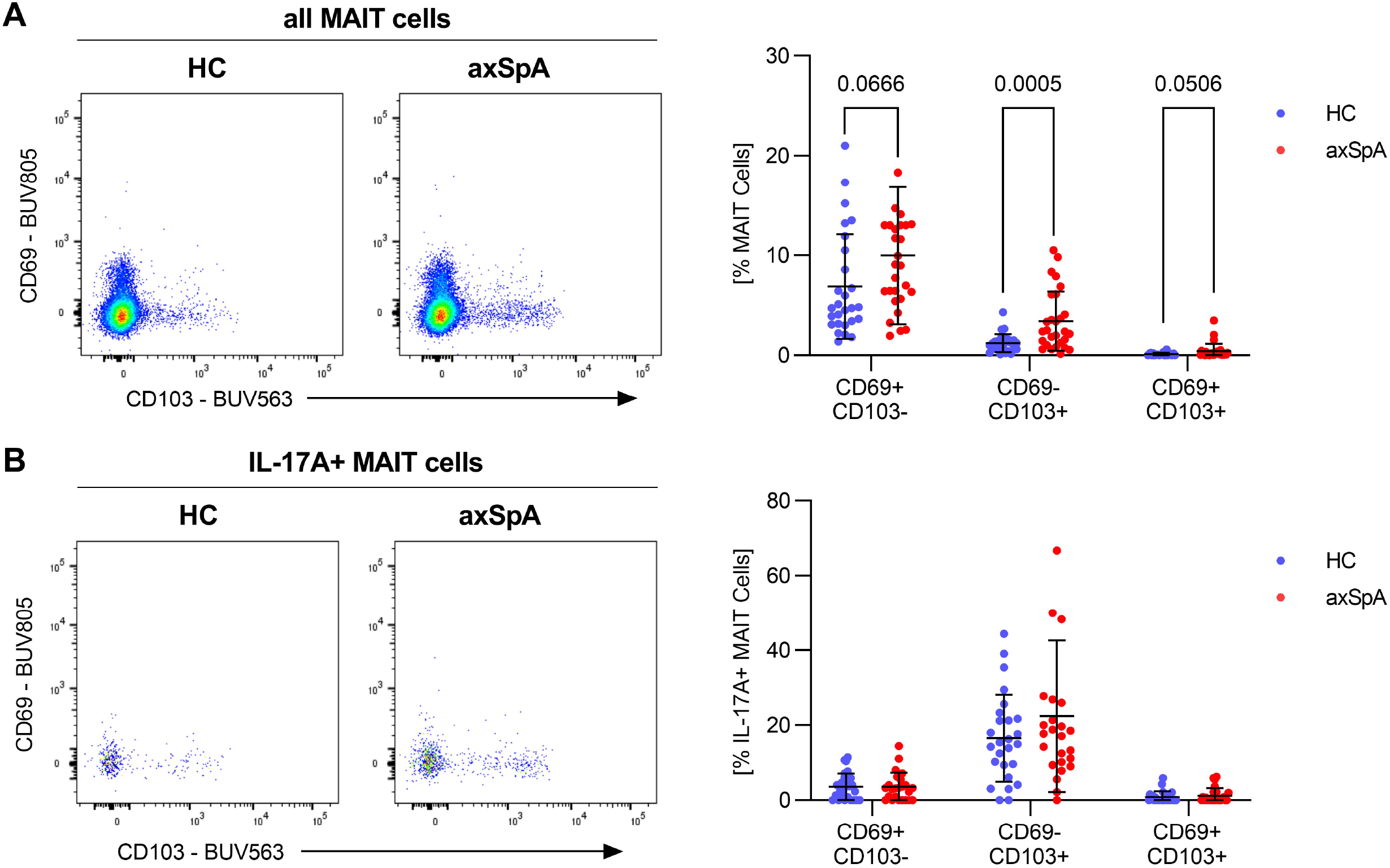
Higher frequency of MAIT cells expressing tissue residency markers in axSpA patients. **A**. MAIT cells from axSpA patients contained a significantly larger fraction of CD69-CD103+ cells (p=0.0005) and with a trend for more CD69+CD103-(p=0.0666) and CD69+CD103+ (p=0.0506) cells compared with healthy controls. **B**. The same analysis for IL-17A+ MAIT cells. CD69-CD103+ MAIT cells had the largest fraction of IL-17A producing cells in both axSpA patients and healthy controls.

**Figure 4.**
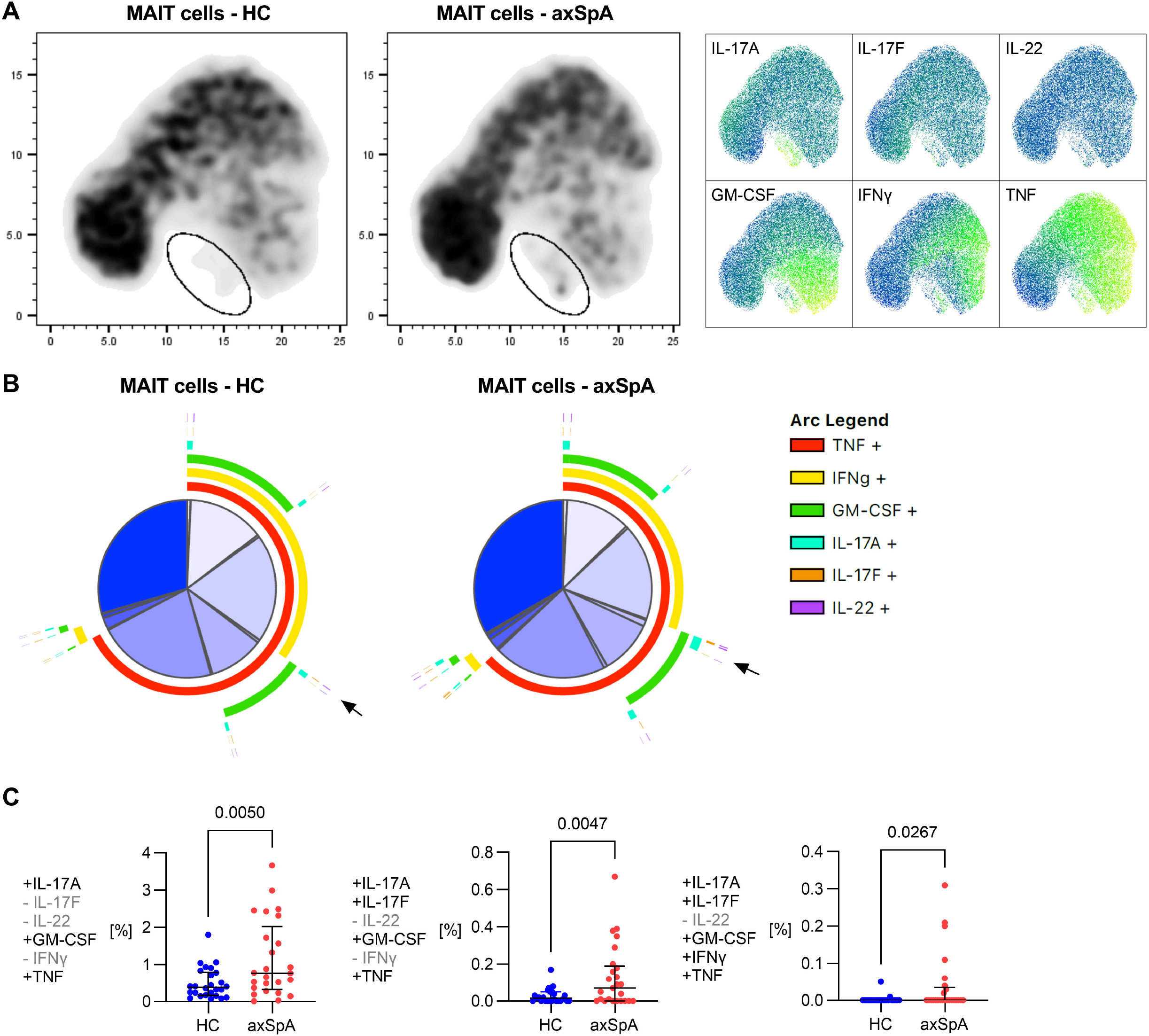
Increased frequency of polyfunctional IL-17A+ MAIT cells in axSpA patients. **A** MAIT cell events from healthy controls and axSpA subjects were concatenated in FlowJo and cytokine expression was analyzed by UMAP. The circled region highlights a cluster of cells with marked difference in abundance between healthy controls and axSpA. Shown on the right are heat maps for each cytokine generated by UMAP clustering of the axSpA samples. The circled region of interest is the primary cluster of IL-17A expressing cells; GM-CSF and TNF are also strongly expressed. **B**. SPICE analysis of the same data. Each slice of the central pie represents the proportion of cells expressing a distinct combination of cytokines. The peripheral arcs show the individual cytokines with overlapping arcs indicating concurrent expression. Black arrows highlight a section of the pie characterized by expression of IL-17A, GM-CSF and TNF that is noticeably bigger in axSpA. **C**. Subject level data show that MAIT cells producing IL-17A, GM-CSF and TNF were expanded in axSpA patients (p=0.0050). MAIT cells that in addition produce IL-17F (p=0.0047) or IL-17F and IFNγ (p=0.0267) were also enriched.

Next, we explored the co-production of cytokines by MAIT cell subsets using a dimension reduction approach. Cytokine expression data for individual MAIT cells from axSpA patients and controls were concatenated and analyzed by UMAP, an algorithm that collapses high-dimensional data into a two-dimensional plot by clustering cells based on similarity of marker expression. Visual inspection of the UMAP plots for axSpA patients and healthy controls identified a small population of cells that was more abundant in patients (Figure 4A). This cluster of cells is characterized by prominent expression of IL-17A with additional expression of IL-17F, GM-CSF and TNF.

**Figure 5.**
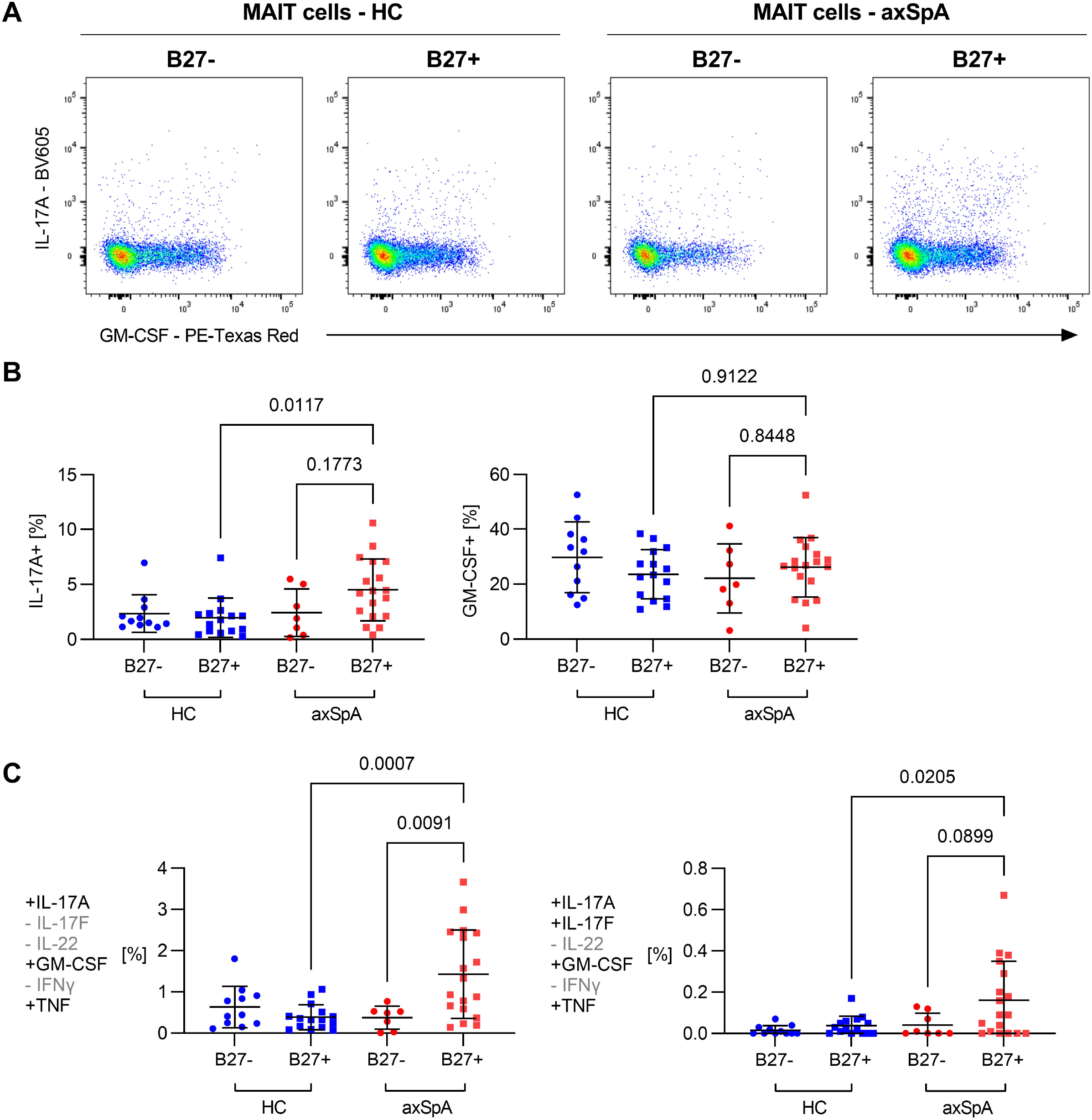
Only HLA-B27+ axSpA patients have expanded IL-17A+ MAIT cells. **A**. Example plots of IL-17A and GM-CSF expression in MAIT cells from healthy controls and axSpA patients that are either HLA-B27+ or HLA-B27-. **B**. MAIT cells from HLA-B27+ axSpA patients, but not HLA-B27-axSpA patients, had a higher fraction of IL-17A+ cells than healthy controls while no difference was detected for GM-CSF expression. **C**. Similarly, MAIT cells expressing IL-17A, GM-CSF and TNF were enriched in HLA-B27+ but not in HLA-B27-axSpA patients (p=0.0007). This relationship holds true for MAIT cells that are positive for IL-17A, IL-17F, GM-CSF and TNF+ (p=0.0205).

We then applied SPICE, an algorithm that was specifically developed to analyze the frequency of cells expressing multiple cytokines in flow cytometry data.[17] The comparison of axSpA and control MAIT cells yielded statistically significant differences in the frequency of 6 populations, all of which included IL-17A as the lead cytokine (Figure 4B). Subject-level data for the populations expressing 3 or more cytokines were extracted from SPICE (Figure 4C): Compared with healthy controls, axSpA patients had higher frequencies of MAIT cells expressing IL-17A, GM-CSF and TNF (p=0.005), IL-17A, GM-CSF, TNF and IL-17F (p=0.0047) and IL-17A, GM-CSF, TNF, IL-17F and IFNγ (p=0.0267).

### MAIT cell in axSpA patients are linked to HLA-B27 status

Lastly, we analyzed whether the expansion of IL-17A+ MAIT cells in axSpA was associated with clinical factors. We observed a non-significant trend toward correlation with CRP but not BASDAI (Suppl. Figure 6A). There was no difference between axSpA patients stratified based on sex or current use of biologic drugs (Suppl. Figure 6B). However, we observed a strong correlation between the frequency of IL-17A+ MAIT cells and HLA-B27 status in axSpA patients, which was not seen in healthy controls (Figure 5B). Polyfunctional IL-17A+ MAIT cell populations were also increased in HLA-B27+ but not in HLA-B27-axSpA patients (Figure 5C).

## Discussion

This study compared peripheral blood lymphocytes in patients with axSpA and healthy controls using 25-parameter fluorescent flow cytometry. Consistent with previous studies, we found an increased frequency of IL-17A+ MAIT cells in axSpA patients even though the total fraction of MAIT cells was reduced.[11–13] IL-17A+ production was enriched in CD69-CD103+ MAIT cells. The IL-17A+ MAIT cell population in axSpA patients contained an increased fraction of polyfunctional cells producing GM-CSF, IL-17F and TNF. The expansion of IL-17A+ MAIT cells was only observed in HLA-B27+ individuals with axSpA.

Prior studies evaluating cytokine expression by peripheral blood lymphocytes in axSpA have yielded discordant results.[5, 6] Multiple factors may explain these discrepancies including differences in the criteria for the selection of patients and controls, variable disease activity between studies or differences in the protocols for in vitro stimulation and staining. The most consistent finding has been a decrease in the fraction of MAIT cells and an increase of IL-17A producing MAIT cells in axSpA.[8, 12, 13] While previous studies used a combination of CD161 and TCRVα7.2 to identify MAIT cells, our study confirms this finding using the more specific MAIT cell definition of MR-1 5-OP-RU tetramer positivity.[25] Together these studies make a strong case that MAIT cells are a major source of IL-17A in axSpA and likely play a role in disease pathogenesis. While we were unable to reproduce other findings of lymphocyte abnormalities reported in the literature, we cannot definitively rule out that additional lymphocyte populations contribute to the exaggerated IL-17A response in axSpA. Larger studies that include more treatment-naive subjects are needed.

A reduction of MAIT cells in peripheral blood is not specific for axSpA and has been described in a variety of settings including rheumatoid arthritis (RA), systemic lupus erythematosus and COVID-19.[26–28] MAIT cells have been found in inflamed non-mucosal tissues including synovial fluid in RA and ankylosing spondylitis.[11, 26, 27] MAIT cells express many integrins and chemokine receptors and are thus poised to move from the blood into inflamed tissues.[29] This could explain their lower peripheral blood numbers in inflammatory diseases. Alternative explanations invoke an increased propensity of MAIT cells for activation-induced cell death and altered exit from mucosal sites.[30] Although MAIT cells are considered to be tissue-resident, they have been shown to be present in human thoracic duct lymph.[31] It is likely that changes in the tissue milieu at epithelial barriers will affect which and how many MAIT cells are allowed to leave, enter the blood stream via the lymphatics and participate in immune surveillance elsewhere in the body. A recent study demonstrated intestinal dysbiosis in HLA-B27+ healthy individuals.[32] Both dysbiosis and intestinal inflammation have been implicated in the pathogenesis of axSpA.[33, 34] Our finding that the expansion of polyfunctional IL-17A+ MAIT cells in the peripheral blood was limited to HLA-B27+ patients may therefore suggest that an HLA-B27 controlled process in the intestine results in the spill-over of primed MAIT cells into the circulation followed by homing into axSpA relevant tissues and local production of pro-inflammatory cytokines. Whether MAIT cells are indeed present in the inflamed pelvis and spine of patients with axSpA has not been reported yet.

We have adopted the term polyfunctional from the work by others [35–37] acknowledging that this may be a suboptimal term as the production of cytokines is arguably a single function. MAIT cells have been shown to express several lineage transcription factors providing them with the capacity to produce multiple cytokines without following the Th1/Th2/Th17 paradigm.[38] Indeed, the pie graphs in Figure 4 demonstrate that MAIT cells in both axSpA patients and healthy controls contain a substantial fraction of cells producing IFNγ, TNF and GM-CSF upon stimulation with PMA/ionomycin. However, the frequency of these polyfunctional MAIT cells was not different between patients and controls. The polyfunctional MAIT cells we found to be expanded in axSpA have IL-17A as the signature cytokine along with variable expression of IL-17F, GM-CSF and TNF. It will be interesting to further explore the heterogeneity of MAIT cells in axSpA under a variety of stimulatory conditions and measuring a higher number of parameters per cell, for instance using single-cell RNA-seq.[21, 39, 40]

Our study has several strengths. Patients and healthy controls were well matched for age, sex and HLA-B27 status. This was achieved by recruiting HLA-B27 genotyped healthy controls through MGB Biobank. The flow cytometry panels permitted the simultaneous analysis of multiple lymphocyte subsets, which had previously only been analyzed in isolation. We further measured 6 cytokines per cell and used MR-1 5-OP-RU tetramers to identify MAIT cells. Limitations include the relatively low disease activity in our axSpA cohort. While the number of subjects included in the analysis was comparable to other studies in this field, more than 60% of axSpA patients were on a biologic and their mean BASDAI score was 2.9, which is relatively low. Inclusion of more biologic-naive patients with active disease would have been desirable but recruitment was difficult given the high frequency of biologic therapy in established axSpA patients in the US.[41, 42] COVID-19 related research restrictions that went into effect in early 2020 added to this problem. Our flow cytometry panels were also not designed to specifically interrogate MAIT cells. In hindsight, inclusion of additional parameters to characterize MAIT cells (e.g. granzymes, CD107, homing receptors) would have been informative.[43]

In conclusion, we demonstrate an expansion of polyfunctional IL-17A+ MAIT cells in the peripheral blood of HLA-B27+ axSpA patients. MAIT cell abnormalities, in particular an increased frequency of IL-17+ MAIT cells, have been the most consistent finding of peripheral blood lymphocyte studies in axSpA to date, suggesting that MAIT cells are a major candidate link between inflammation in the gut and inflammation in the axial skeleton.

## Acknowledgements

We would like to thank Dr. Lianne Gensler, UCSF, for providing the DX31 antibody. Jake Lawicky, Greg Keras, H. Maura Friedlander, Alessandra Zaccardelli, Lauren Prisco and Lily Marin helped with patient recruitment.

