## Supplement for "Polyfunctional IL-17A+ MAIT cells are expanded in the peripheral blood of patients with HLA-B27+ axial spondyloarthritis"

**Supplemental Table 1. Markers and antibodies – staining panel I.**

| Laser | Flurochrome | Marker | Clone | Company |
|---|---|---|---|---|
| **355 nm** | eFluor 455UV | FVD | | Thermo Fisher Scientific |
| | BUV496 | CD8a | RPA-T8 | BD Bioscience |
| | BUV563 | CD103 | Ber-ACT8 | BD Bioscience |
| | BUV615 | CD49a | SR84 | BD Bioscience |
| | BUV661 | PD-1 | EH12.1 | BD Bioscience |
| | BUV737 | CD14 | M5E2 | BD Bioscience |
| | BUV805 | CD69 | FN50 | BD Bioscience |
| **405 nm** | BV421 | MR-1 Tetramer | 5-OP-RU loaded | NIH |
| | BV510 | CD4 | OKT4 | Biolegend |
| | BV570 | CD45RA | HI100 | Biolegend |
| | BV605 | CCR7 | G043H7 | Biolegend |
| | BV650 | CD25 | BC96 | Biolegend |
| | BV711 | vδ2 TCR | B6 | Biolegend |
| | BV750 | αβ TCR | IP26 | Biolegend |
| | BV786 | CD161 | HP-3G10 | Biolegend |
| **488 nm** | FITC | vδ1 TCR | REA173 | Miltenyi Biotec |
| | PerCP-Cy5.5 | CD16 | 3G8 | Biolegend |
| | | CD203c | NP4D6 | Biolegend |
| | | FcεRI | AER-37 (CRA-1) | Biolegend |
| | | CD34 | 561 | Biolegend |
| **561 nm** | PE | CD56 | 5.1H11 | Biolegend |
| | PE-Texas Red | CD127 | A019D5 | Biolegend |
| | PE-Cy5 | CD19 | HIB19 | Biolegend |
| | PE-Cy7 | KIR3DL1 | DX9 | Biolegend |
| **637 nm** | APC | KIR3DL2,<br>anti-mouse IgG | DX31,<br>polyclonal | UCSF Monoclonal Core,<br>Jackson ImmunoResearch |
| | Alexa Fluor 700 | CD3 | UCHT1 | Biolegend |
| | APC-Cy7 | C-KIT | 104D2 | Biolegend |

**Supplemental Table 2. Markers and antibodies – staining panel II.**

| Laser | Flurochrome | Marker | Clone | Company |
|---|---|---|---|---|
| **355 nm** | eFluor 455UV | FVD | | Thermo Fisher Scientific |
| | BUV496 | CD8a | RPA-T8 | BD Bioscience |
| | BUV563 | CD103 | Ber-ACT8 | BD Bioscience |
| | BUV615 | CD49a | SR84 | BD Bioscience |
| | BUV661 | PD-1 | EH12.1 | BD Bioscience |
| | BUV737 | IFNγ | 4S.B3 | BD Bioscience |
| | BUV805 | CD69 | FN50 | BD Bioscience |
| **405 nm** | BV421 | MR-1 Tetramer | 5-OP-RU loaded | NIH |
| | BV510 | CD4 | OKT4 | Biolegend |
| | BV570 | CD45RA | HI100 | Biolegend |
| | BV605 | IL-17A | BL168 | Biolegend |
| | BV650 | IL-17F | O33-782 | BD Bioscience |
| | BV711 | vδ2 TCR | B6 | Biolegend |
| | BV750 | αβ TCR | IP26 | Biolegend |
| | BV786 | CD161 | HP-3G10 | Biolegend |
| **488 nm** | FITC | vδ1 TCR | REA173 | Miltenyi Biotec |
| | PerCP-Cy5.5 | KIR3DL1 | DX9 | Biolegend |
| **561 nm** | PE | CD56 | 5.1H11 | Biolegend |
| | PE-Texas Red | GM-CSF | BVD2-21C11 | Biolegend |
| | PE-Cy5 | CD14 | 61D3 | Thermo Fisher Scientific |
| | | CD19 | HIB19 | Biolegend |
| | | CD33 | WM53 | Biolegend |
| | PE-Cy7 | IL-22 | 2G12A41 | Biolegend |
| **637 nm** | APC | KIR3DL2, anti-mouse IgG | DX31, polyclonal | UCSF Monoclonal Core, Jackson ImmunoResearch |
| | Alexa Fluor 700 | CD3 | UCHT1 | Biolegend |
| | APC-Cy7 | TNF | Mab11 | Biolegend |

**Supplemental Figure 1. Flow diagrams.**

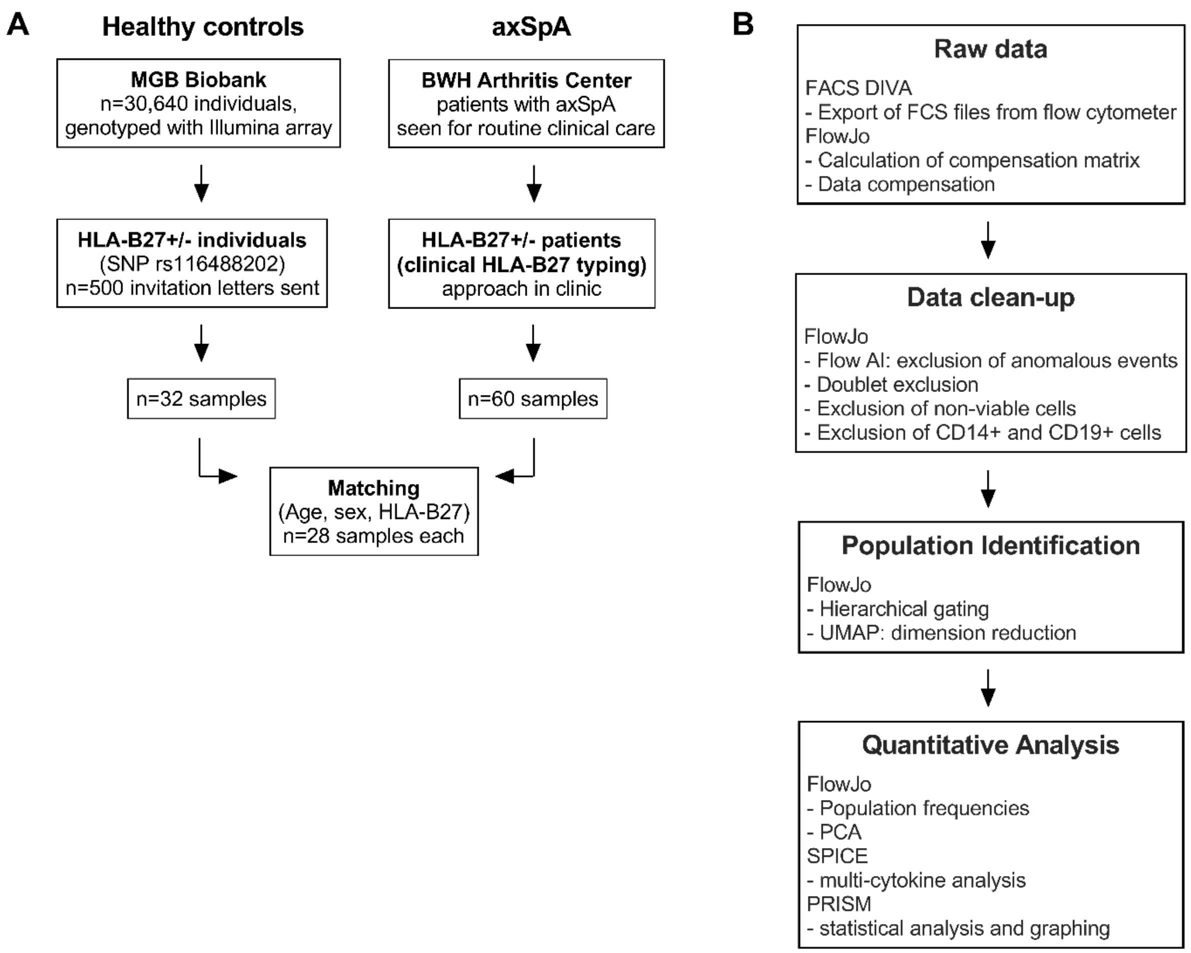


**A.** Subject recruitment. Healthy controls were recruited through MGB Biobank. Patients with axSpA were recruited in the BWH arthritis center. n=28 samples from each group matched for age, sex and HLA-B27 status were analyzed. **B.** Data analysis. Data were collected on a FACSymphony flow cytometer. Uncompensated raw data were exported as FCS files and imported into FlowJo. Details of data cleanup, populations identification and quantitative analysis are provided in Supplemental Methods.

**Supplemental Figure 2. Gating strategy.**

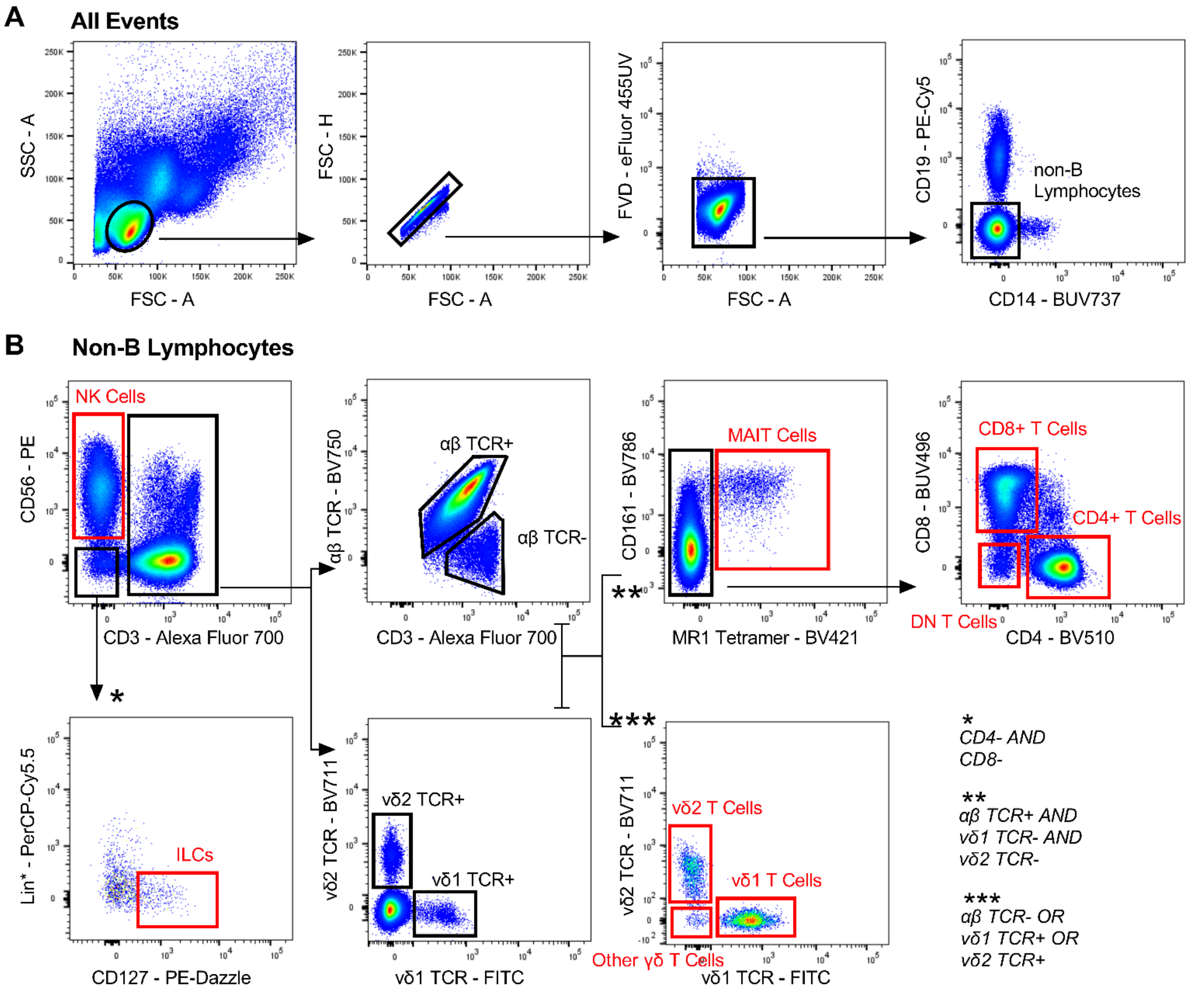


Representative plots from an axSpA sample demonstrating the hierarchal gating strategy for major lymphocyte populations in peripheral blood. **A.** Lymphocytes were selected according to forward and side scatter, followed by sequential exclusion of doublets, FVD positive non-viable cells, CD19+ B cells and CD14+ monocytes. The population of CD19-CD14- cells (non-B lymphocytes) in the plot on the very right represents the reference population for the quantification of lymphocyte populations. **B.** NK cells were identified as CD56+ and CD3-. Boolean gating was used to differentiate CD3+ cells into αβ and γδ T cells. αβ T cells were identified as αβ TCR+ AND Vδ1 TCR- AND Vδ2 TCR-. γδ T cells were identified as αβ TCR- OR Vδ1 TCR+ OR Vδ2 TCR+. In the αβ T cell compartment, MAIT cells were gated as MR-1 5-OP-RU tetramer positive. MR-1 5-OP-RU tetramer negative cells were subdivided into CD4+, CD8+, and double negative αβ T cells. In the γδ T cell compartment, cells were subdivided into Vδ1+, Vδ2+, and other γδ T cells. In the CD3-CD56- gate, ILCs were identified by first excluding CD4 and CD8 and then selecting CD127+ and lineage (CD16, CD34, CD203c, FcεRI) negative events.

**Supplemental Figure 3. Identification of CD4+ and CD8+ T cell subsets.**

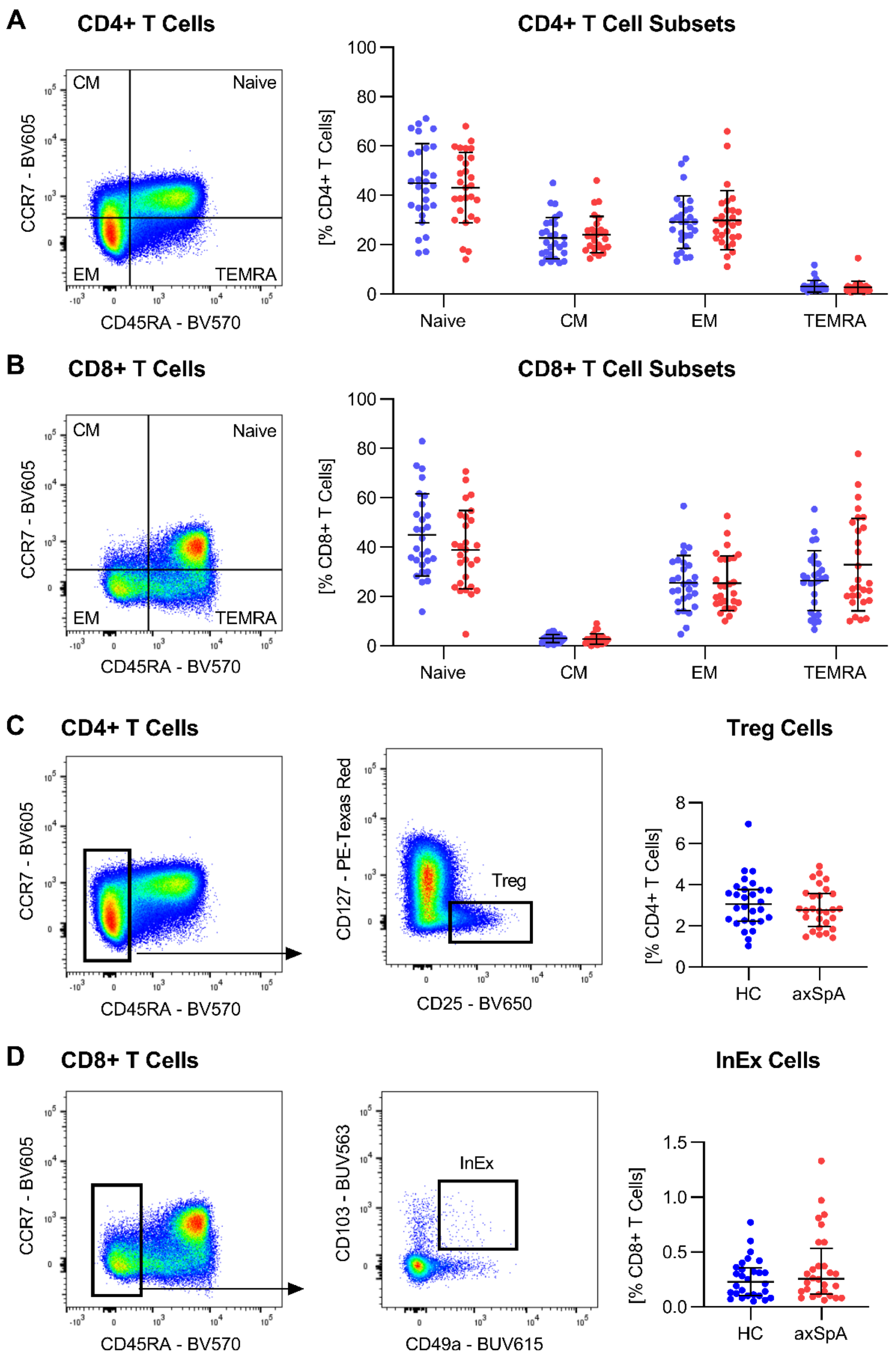


**A.** Using CCR7 and CD45RA, CD4+ T cells were divided into naïve (CD45RA+CCR7+), central memory (CM, CD45RA-CCR7+), effector/memory (EM, CD45RA-CCR7-) and terminally differentiated effector/memory (TEMRA, CD45RA+CCR7-) subsets. **B.** Subsets of CD8+ T cells were similarly identified using CCR7 and CD45RA. **C.** Treg cells were identified within the CD4+ T cell gate as

CD45RA-CD127-CD25+. **D.** InEx cells were identified within the CD8+ T cell gate as CD45RA-CD103+CD49a+. There were no statistically significant differences between axSpA patients and healthy controls for any of these T cell subsets.

**Supplemental Figure 4. Cytokine expression in major lymphocyte populations stimulated with PMA/ionomycin.**

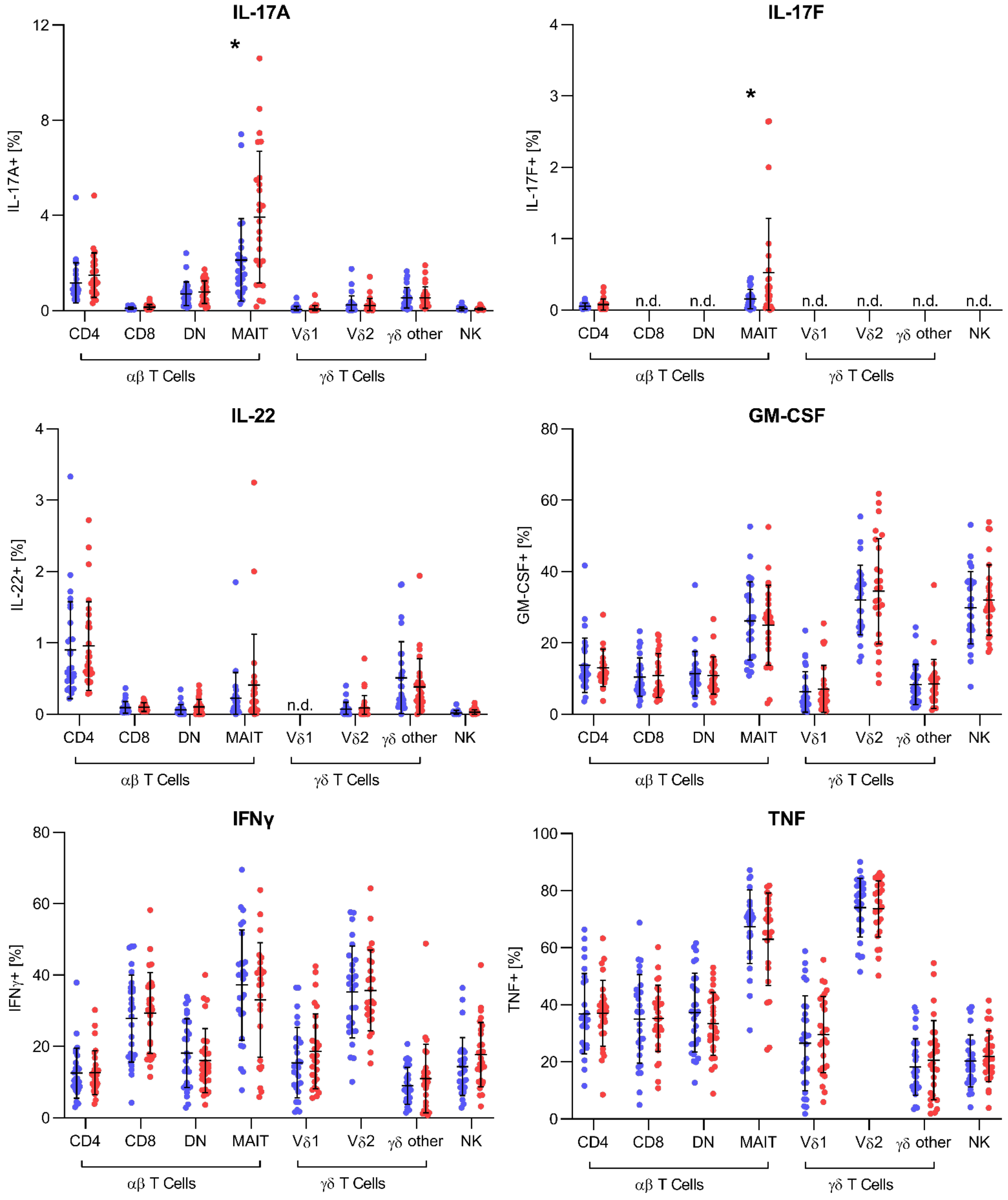


The fraction of positive cells for each of the 6 measured cytokines in 9 major lymphocyte populations plots. An asterisk indicates a p value < 0.05. The only statistically significant difference between axSpA patients and healthy controls was seen in IL-17A+ MAIT cells and IL-17F+ MAIT cells.

**Supplemental Figure 5. Characterization of MAIT cells.**

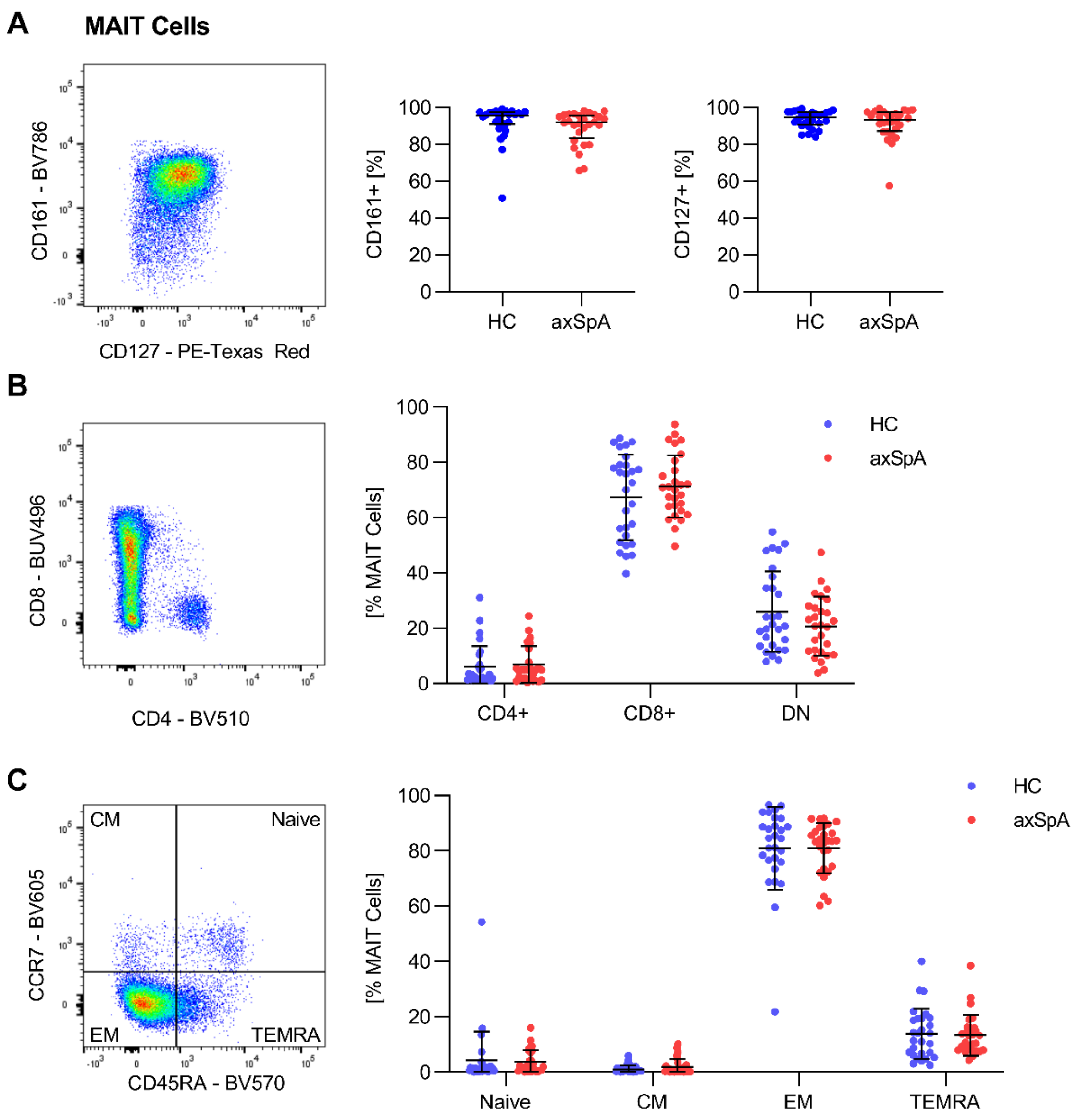


**A.** Example plot and quantification of CD127+ and CD161+ MAIT cells. No difference between axSpA patients and healthy controls. **B.** Example plot and quantification of CD4+ and CD8+ MAIT cells. No difference between axSpA patients and healthy controls. **C.** Subsets of MAIT cells defined by expression of CCR7 and CD45RA. There was no difference between axSpA patients and healthy controls.

**Supplemental Figure 6. Correlation of MAIT cell IL-17A expression with clinical parameters.**

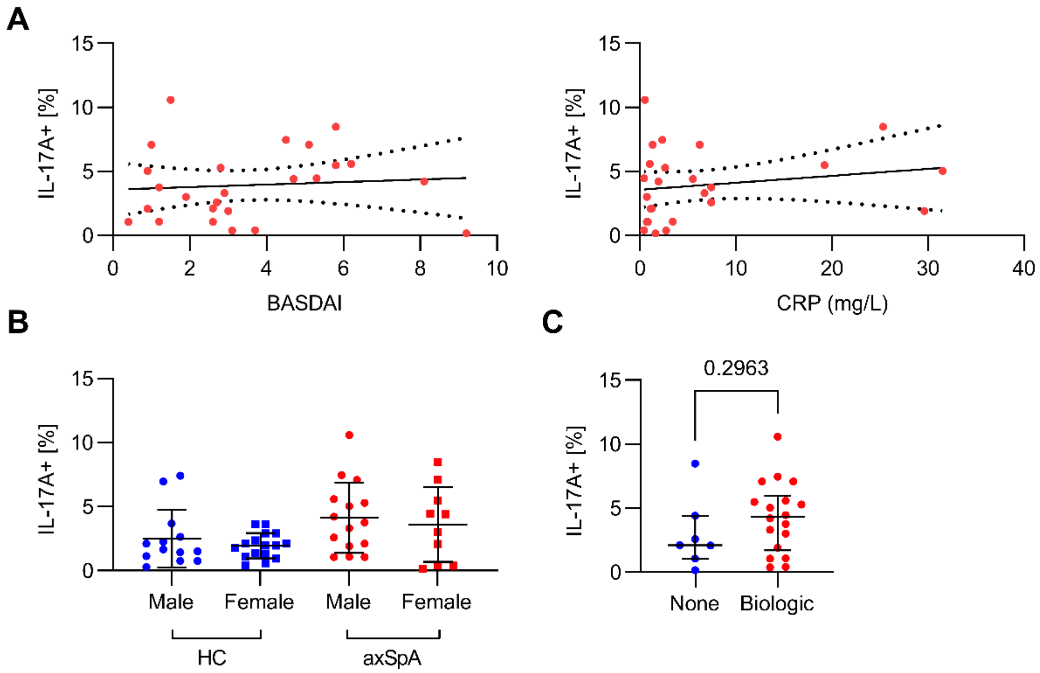


**A.** The fraction of IL-17A+ MAIT cells plotted against BASDAI, no significant trend was observed. IL-17A+ MAIT were also cells plotted against serum CRP. There were statistically non-significant trends for increased IL-17A expression with increased inflammatory activity. **B.** No difference in the fraction of IL-17A+ MAIT cells between male and female patients with axSpA. **C.** There was no difference in the fraction of IL-17A+ MAIT cells between axSpA patients taking or not taking a biologic DMARD.

**Supplemental Methods**

**Antibody Staining**

Cells were washed with Hanks' Balanced Salt Solution (HBSS) (Corning) and viability staining was performed using Fixable Viability Dye (FVD) eFluor 455UV (Thermo Fisher Scientific). Fc receptors were blocked using Human TrueStain FCX (BioLegend). Cells were stained at room temperature with MR-1 5-OP-RU tetramer (NIH Tetramer Core Facility) and monoclonal anti-KIR3DL2 antibody (clone DX31, UCSF Monoclonal Antibody Core Facility). Cells were washed and stained with APC-labelled anti-mouse IgG secondary antibody (Jackson ImmunoResearch). Cells were washed again and stained for all other surface markers for 30 minutes at 4° C. Stimulated cells were then fixed using IC Fixation Buffer (Thermo Fisher Scientific) and intracellular staining was done in Permeabilization Buffer (Thermo Fisher Scientific) for 30 minutes at room temperature in the dark.

**Compensation and data clean-up**

Cytometry data were collected on a FACSymphony flow cytometer (BD Bioscience). Raw data were exported as FCS files. Compensation matrix was calculated using FlowJo followed by manual adjustments where needed. Data clean-up was performed in FlowJo v10.8.0 using FlowAI v2.2 to remove anomalous events.[1] Doublets were excluded based on relationship of forward scatter area and height. Cells that stained positive with FVD were excluded as non-viable. Monocytes and B cells were excluded based on expression of CD14 and CD19 respectively. Lymphocyte populations were identified either by hierarchal gating or by dimensional reduction with UMAP v3.1.[2] PCA was used to examine global differences in lymphocyte subset frequency between healthy controls and axSpA.

**FACS Plot Generation**

Example FACS plots of MAIT cells in Figures 3 and 5 were created by combining FCS files from either the healthy controls or axSpA patients. First, all FCS files were down-sampled to 1000 events. Subjects with fewer than 500 events were excluded (2 healthy controls and 3 axSpA). The remaining FCS files were concatenated into either a healthy control file or axSpA file. The healthy control file contained a greater number of events, so it was further down-sampled to match the number of the axSpA events.

**PCA**

Population frequencies from each sample were imported into R and PCA was performed using the prcomp function.[3] Visualization of PCA data was done with ggplot2.[4] Ellipses in Figure 1 represent a 95% confidence interval.

**UMAP**

The number of MAIT events from all healthy controls and axSpA subjects were down-sampled in FlowJo to 1000 per subject. Subjects with fewer than 500 events were excluded to limit error due to mismatched sample weighting. Two healthy control and three axSpA samples were removed. Healthy control and axSpA events were concatenated into two separate files. Down-sampling was used again to equal weight the contribution from both subject sets. The balanced healthy control and axSpA files were then concatenated into a single FCS file which was analyzed using the FlowJo UMAP module.

**SPICE**

Subjects with fewer than 500 MAIT cell events were excluded from cytokine analysis. In FlowJo, Boolean gating was used to identify all possible cytokine combinations. As each cell can be either negative or positive for a single cytokine, a combination of 6 cytokine antibodies yields 64 possible groups ranging from expressing not a single cytokine to being positive for all 6 cytokines. Data were then exported into SPICE for analysis and visualization.[5] In the SPICE plot, the 64 groups are arranged as pie slices in the center, the production of the 6 cytokines for each of the 64 groups are shown as peripheral, overlapping arcs.
